# The dual role of the angiosperm radiation on insect diversification

**DOI:** 10.1101/2023.02.07.527317

**Authors:** David Peris, Fabien L. Condamine

**Affiliations:** Institut Botànic de Barcelona (CSIC-Ajuntament de Barcelona), 08038 Barcelona, Spain; CNRS, UMR 5554 Institut des Sciences de l’Evolution de Montpellier, Université de Montpellier, Place Eugène Bataillon, 34095 Montpellier, France

## Abstract

Most of the animal pollination is realized by insects, interactions between them and flowering plants have been hypothesized to be important drivers of diversification. Yet, there is weak support for coevolutionary diversification in plant–pollinator interactions. Macroevolutionary studies on insect and plant diversities support the hypothesis that angiosperms evolved after an insect diversity peak in the Early Cretaceous, suggesting that gymnosperm pollinators may have been accessible for angiosperms when they evolved. We examined fossil and phylogenetic evidence documenting this hypothesis and provide new clues on the impact of angiosperm radiation on insect diversification. Using the family-level fossil record of insects and a Bayesian process-based approach, we estimated diversification rates and the role of six different variables on insect macroevolutionary history. We found that, among the six tested variables, angiosperms had a dual role that has changed through time with an attenuation of insect extinction in the Cretaceous and a driver of insect origination in the Cenozoic. However, increasing insect diversity, spore plants and global temperature also showed strong positive correlation with both origination and extinction rates of insects, suggesting that different drivers had important effect on insect evolution, not just angiosperms, which would deserve further studies.

## Introduction

Pollination is an essential activity in the sexual reproduction of many plants, particularly for plants producing seeds. The difficulty of active movement in most plants makes necessary the action of different pollination agents to transport their male gametes contained within pollen to ovule bearing, female structures of the same species. These transport agents can be wind, water, or vertebrate animals, although most plants require principally the action of invertebrate animals for their reproduction (Ollerton, 2017). Most current pollinators belong to one of the four major insect orders: Hymenoptera, Diptera, Lepidoptera and Coleoptera (Ollerton, 2017). Iconic pollinator groups currently include bees and butterflies. However, a variety of seed plants existed and possessed reproductive organs long before the evolution of these more notable pollinator groups. This suggests that ancient plants were pollinated by a different spectrum of pollinator agents (Labandeira, 2007; Peris et al., 2017; Khramov et al., 2020).

Gymnosperms dominated the land surface until their sister clade, the angiosperms, experienced a rapid diversification in the Cretaceous, for some authors called the Angiosperm Radiation (125–90 Ma, *e*.*g*., Labandeira, 2014), the Cretaceous Terrestrial Revolution (125–80 Ma, Lloyd et al., 2008), but recently also known as the Angiosperm Terrestrial Revolution (ATR, 100–50 Ma, Benton et al., 2022) (Figure 1). This turnover likely caused a decline of conifers (Condamine et al., 2020) and other plant lineages (Lidgard and Crane, 1988). First plants were wind pollinated until some insects diversified and started to feed on gymnosperm ovular secretions in a surface-fluid-feeding way or gymnosperm pollen (Labandeira 1998), predating that of nectar-feeding insects on angiosperms (Nepi et al., 2017) (Figure 1).

**Figure 1.**
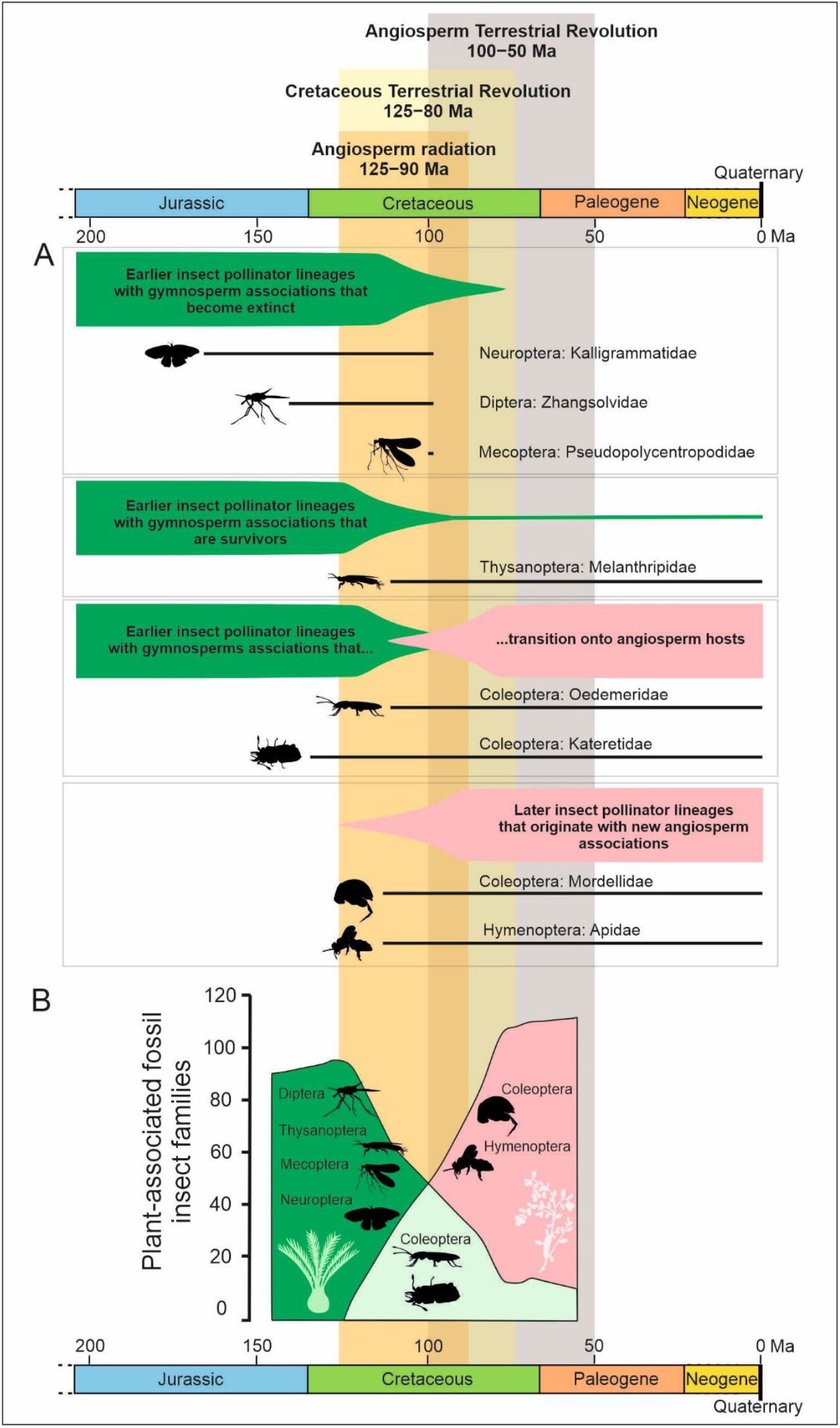
Different Cretaceous pollination modes described in Peris et al. (2017), showing different groups of insects under distinctive patterns of extinction, survival, and origination following the gymnosperm-angiosperm transition. The fossil pollination cases reviewed in Peris et al. (2020) and Peña-Kaitath et al. (in prep.). The period of angiosperm evolution is marked as Angiosperm Radiation (Labandeira, 2014), Cretaceous Terrestrial Revolution (Lloyd et al., 2008), and Angiosperm Terrestrial Revolution (Benton et al., 2022). **A**. Diverse fossil community of Cretaceous pollinators and the lifespan of these families. **B**. Representation of the transition from gymnosperm–insect pollination to angiosperm–insect pollination with the transitional examples.

The complex interactions between pollinators and gymnosperms seems to have been persisting since, at least, the early Permian (283–273 million years ago, Ma), predating the first flowering plants by more than 100 Ma (Khramov et al., 2022). There is a diverse well-documented herbivore community from the Cretaceous found in sediments and ambers supporting gymnosperm–insect pollination modes and host associations with ginkogaleans, cycads, conifers and bennettitalean gymnosperms during the Early and the beginning of the Late Cretaceous (Labandeira, 2007; Peris et al., 2017, 2020; Khramov et al., 2020) (Figure 1). By contrast, evidence for angiosperm-pollen consumers or flower visitors appeared only in the Late Cretaceous, around 99 Ma (Peris et al., 2020; Peña-Kaitath et al., in prep.). It is thought that flowering plants have interacted with pollinating insects since their beginning (Grimaldi, 1999; Labandeira, 2007), but gymnosperm-adapted insects were not using extensively angiosperms resources yet until the Late Cretaceous (Labandeira, 2007; Peris et al., 2017; 2020). Elucidating when, how and why the ecological transformation of ecosystems began with the coevolution between insects and flowering plants has become an idea of recent interest (van der Kooi and Ollerton, 2020; Benton et al., 2022; Asar et al., 2022).

The great radiation of modern insect lineages started 245 Ma (Labandeira and Sepkoski, 1993; Rainford et al. 2014). Since the Jurassic, insect families showed low extinction rates (Labandeira and Sepkoski, 1993; Jarzembowski and Ross, 1996; Condamine et al. 2016). Insect family-richness peaked during the Early Cretaceous around 125 Ma, when angiosperms were still rare (Claphman et al., 2016; Schachat et al., 2019). This peak occurred prior to extinctions within early-diverging groups, in part linked to mid-Cretaceous floral turnover following the evolution of flowering plants (Claphman et al., 2016; Condamine et al., 2020) (Figure 2). The Early Cretaceous richness peak may therefore reflect a transitional period in insect evolution where radiating extant families coexisted with early-diverging ones that are rare today or that became extinct (Peris et al., 2014; 2020b; Nel et al., 2015; Claphman et al., 2016). The overlap between insects that had specialized relationships with gymnosperms and insects that had specialized relationships with angiosperms may have also contributed to the Early Cretaceous peak in insect diversity (Peris et al., 2017; 2020b; Schachat et al., 2019).

**Figure 2.**
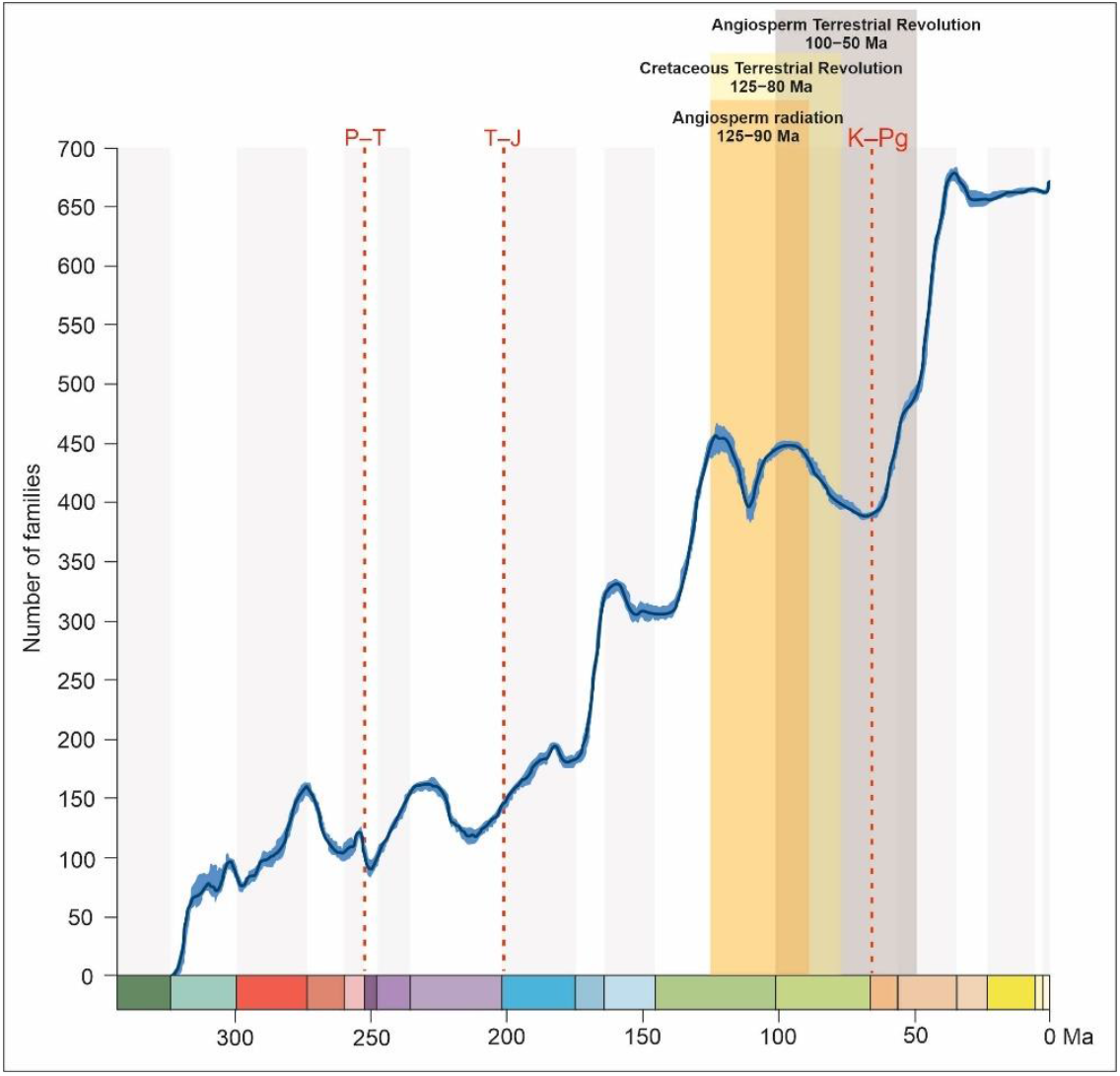
Insect family accumulation through time using the software PyRate and the family-level fossil dataset of Condamine et al. (2016). Red dot lines indicate the Permian–Triassic (P–T), Triassic–Jurassic (T–J), and the Cretaceous–Paleogene (K–Pg) mass extinction. The period of angiosperm evolution is marked as Angiosperm Radiation (Labandeira, 2014), Cretaceous Terrestrial Revolution (Lloyd et al., 2008), and Angiosperm Terrestrial Revolution (Benton et al., 2022).

Here we reviewed the evidence on the origin of flowering plants and their related pollinator insect lineages, and we tested the impact of angiosperm radiation on insect diversification. We analysed the role that six different variables (diversity dependence, angiosperms, gymnosperms, spore plants, continental fragmentation, and temperature) have on diversification rates of insects (Figure 3).

**Figure 3.**
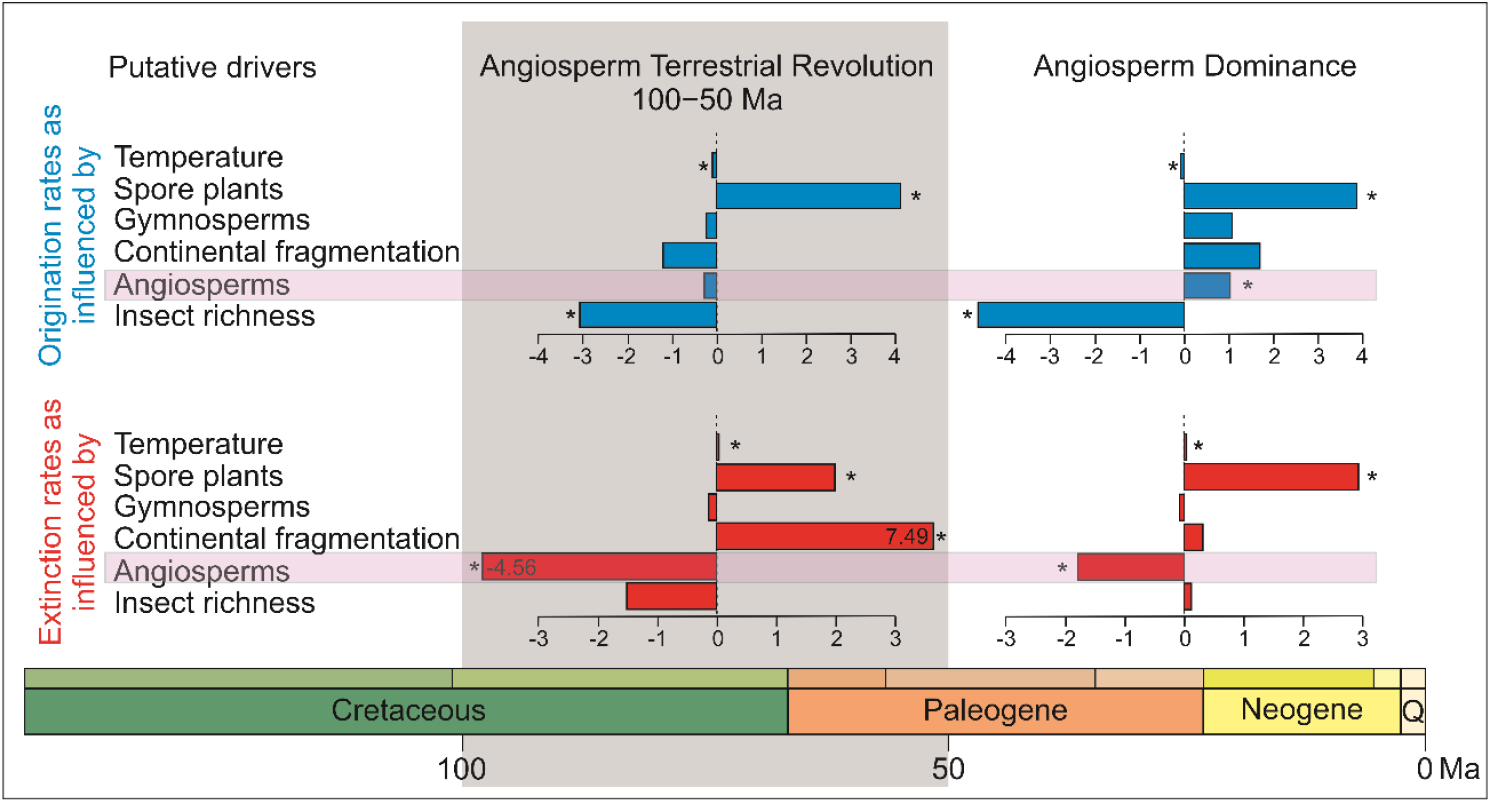
Correlation trends of different analysed drivers for origination (in blue) and extinction (in red) rates on insect diversity for the Angiosperm Terrestrial Revolution timeframe (100–50 Ma, Benton et al., 2022) and for the Angiosperm Dominance period (50–0 Ma). Data used in this representation is offered in the Table 1. Drivers from top to bottom are Global mean temperature (Temperature), Spore plant relative diversity (Spore plants), Gymnosperm relative diversity (Gymnosperms), Continental fragmentation, Angiosperm relative diversity (Angiosperms), and Insect family richness (Insect richness). Asterisks indicate significant correlations recovered with the MBD model (shrinkage weight >0.5 and 95% CI not overlapping with zero).

### Evolution of angiosperms

Today angiosperms dominate most terrestrial ecosystems and provide an important part of the food chain and niche requirements for many other organisms. Estimations indicate that flowering plants represent about 90% of all land plants (embryophytes), which is around 300,000 living species (Christenhusz and Byng, 2016). The present-day diversity of species and forms in flowering plants have intrigued biologists for more than 150 years (Buggs, 2021). Their origin was one of the most transformative events in Earth history (Figures 1–2). However, the age of crown angiosperms remains highly uncertain (Sauquet et al., 2022), despite recent claims of more definite answer to when exactly angiosperms originated and began to diversify (*e*.*g*., Barba-Montoya et al., 2018; Li et al., 2019; Silvestro et al., 2021). The stem age of angiosperms, their divergence time from extant gymnosperms, is dated as 310–380 Ma depending on the study (Clarke et al., 2011; Magallón et al., 2013, Silvestro et al., 2015, Nie et al., 2020). By contrast, the morphological age of flowering plants, that is the age when they are morphology identifiable in the fossil record, is not yet well understood, variously dated from 250–140 Ma (Sauquet and Magallón, 2018).

**Table 1.** Results of the multivariate birth-death model applied to insects in three set of analyses. The table reports the mean and median posterior parameter estimates (and 95% credibility interval, CI) for the parameters of the MBD model: baseline speciation (λ0), extinction rates (μ0) and correlation parameters (Gλ and Gμ) for each of the environmental drivers. Shrinkage weights (ω) greater than 0.5 (highlighted in bold) indicating significant evidence for correlation (positive or negative depending on the respective Gλ or Gμ value).

|  | a) Global insect diversification |  |  | b) Diversification during the ATR (100 to 50 Ma) |  |  | c) Diversification after the ATR (50 Ma to present) |  |  |
| --- | --- | --- | --- | --- | --- | --- | --- | --- | --- |
| Parameters | Mean | Median | 95% CI | Mean | Median | 95% CI | Mean | Median | 95% CI |
| Baseline origination rate $\lambda_0$ | 0.4907 | 0.3092 | [0.1419, 1.196] | 0.8383 | 0.3012 | [0.0754, 3.278] | 0.4248 | 0.1669 | [0.0206, 1.0825] |
| Baseline extinction rate $\mu_0$ | 4.8284E-3 | 4.5032E-3 | [1.9721E-3, 8.4722E-3] | 7.542E-3 | 6.6918E-3 | [2.2163E-3, 0.0149] | 4.7067E-3 | 4.4166E-3 | [1.9018E-3, 7.9883E-3] |
| G $\lambda$ Insect diversity | **-5.0235** | **-4.9936** | **[-5.9243, -4.1341]** | **-3.0585** | **-3.0844** | **[-5.1351, -0.8334]** | **-4.6166** | **-4.6026** | **[-5.9835, -2.983]** |
| G $\lambda$ Angiosperms | **0.8165** | **0.8283** | **[0.1442, 1.4328]** | -0.3412 | -0.2878 | [-1.3542, 0.5499] | **1.0401** | **1.0524** | **[0.1234, 1.9712]** |
| G $\lambda$ Continental frag. | 2.1159 | 2.1251 | [0.7511, 3.453] | -1.3638 | -1.216 | [-5.057, 2.2001] | 1.695 | 1.7441 | [-0.1398, 3.392] |
| G $\lambda$ Gymnosperms | 0.7459 | 0.7819 | [-0.0773, 1.5622] | -0.3112 | -0.2339 | [-1.29, 0.4962] | 1.0731 | 1.1089 | [-0.0532, 2.3286] |
| G $\lambda$ Spore plants | **3.6681** | **3.6892** | **[2.9137, 4.3556]** | **4.0704** | **4.108** | **[3.0406, 4.9928]** | **3.8781** | **3.8383** | **[3.0373, 4.7885]** |
| G $\lambda$ Temperature | **-0.0822** | **-0.0807** | **[-0.1097, -0.0564]** | **-0.0935** | **-0.0925** | **[-0.1368, -0.0469]** | **-0.0683** | **-0.0643** | **[-0.1125, -0.0159]** |
| G $\mu$ Insect diversity | 0.1455 | 0.0932 | [-0.524, 0.9017] | -1.4949 | -1.5064 | [-3.1339, 0.2498] | 0.164 | 0.1091 | [-0.5198, 0.9159] |
| G $\mu$ Angiosperms | **-1.8155** | **-1.7999** | **[-2.4776, -1.1818]** | **-4.5855** | **-4.5552** | **[-6.4712, -2.7859]** | **-1.8073** | **-1.7907** | **[-2.4664, -1.1584]** |
| G $\mu$ Continental frag. | 0.4273 | 0.3555 | [-0.8756, 2.0666] | **7.4912** | **7.486** | **[2.1141, 12.8511]** | 0.4049 | 0.313 | [-0.8913, 2.0322] |
| G $\mu$ Gymnosperms | -0.1271 | -0.0763 | [-0.805, 0.5246] | -0.1786 | -0.1251 | [-0.9781, 0.4926] | -0.117 | -0.0729 | [-0.8323, 0.4772] |
| G $\mu$ Spore plants | **2.887** | **2.8952** | **[2.1342, 3.6284]** | **1.9783** | **1.9942** | **[0.9419, 2.9952]** | **2.9128** | **2.921** | **[2.2031, 3.6651]** |
| G $\mu$ Temperature | **0.0335** | **0.0337** | **[0.0179, 0.0489]** | **0.0315** | **0.0318** | **[0.0144, 0.049]** | **0.0338** | **0.0339** | **[0.0183, 0.0487]** |
| $\omega$ $\lambda$ Insect diversity | **0.9381** | **0.9551** | **[0.8172, 1]** | **0.8511** | **0.9029** | **[0.5166, 1]** | **0.9248** | **0.9472** | **[0.773, 1]** |
| $\omega$ $\lambda$ Angiosperms | **0.5585** | **0.5763** | **[0.0881, 0.9999]** | 0.4305 | 0.392 | [2.9613E-8, 0.9566] | **0.6025** | **0.6449** | **[0.0957, 0.9997]** |
| $\omega$ $\lambda$ Continental frag. | 0.4579 | 0.4197 | [0.0435, 0.9816] | 0.4203 | 0.3781 | [1.0981E-7, 0.9498] | 0.4007 | 0.3402 | [3.0291E-7, 0.9413] |
| ω λ Gymnosperms | 0.4432 | 0.405 | [2.4415E-7, 0.9547] | 0.3536 | 0.2545 | [2.7701E-8, 0.941] | 0.4942 | 0.507 | [1.1021E-8, 0.9625] |
| ω λ Spore plants | **0.861** | **0.8913** | **[0.6245, 1]** | **0.8851** | **0.9139** | **[0.6791, 1]** | **0.8712** | **0.9** | **[0.6482, 1]** |
| ω λ Temperature | **0.8783** | **0.9098** | **[0.6531, 1]** | **0.8978** | **0.9309** | **[0.6817, 1]** | **0.801** | **0.8729** | **[0.315, 1]** |
| ω μ Insect diversity | 0.3319 | 0.2344 | [3.4958E-9, 0.9201] | 0.6806 | 0.7705 | [0.0725, 1] | 0.3418 | 0.2428 | [1.183E-7, 0.9353] |
| ω μ Angiosperms | **0.7614** | **0.8017** | **[0.4099, 0.9999]** | **0.9264** | **0.9502** | **[0.7781, 1]** | **0.7592** | **0.7968** | **[0.4084, 1]** |
| ω μ Continental frag. | 0.2669 | 0.136 | [1.7915E-8, 0.8964] | **0.7439** | **0.8034** | **[0.2966, 1]** | 0.2667 | 0.1316 | [7.8828E-12, 0.8947] |
| ω μ Gymnosperms | 0.2797 | 0.1553 | [3.1608E-8, 0.9049] | 0.3161 | 0.1968 | [2.8001E-10, 0.9324] | 0.2744 | 0.1524 | [5.2828E-8, 0.8987] |
| ω μ Spore plants | **0.8145** | **0.851** | **[0.5176, 1]** | **0.7404** | **0.7845** | **[0.348, 1]** | **0.8195** | **0.8563** | **[0.5299, 1]** |
| ω μ Temperature | **0.6892** | **0.7259** | **[0.2751, 1]** | **0.6926** | **0.7342** | **[0.2646, 0.9998]** | **0.6846** | **0.7215** | **[0.2798, 0.9998]** |

The earliest fossil remains that can be assigned with high confidence to crown-group angiosperms are tricolpate pollen grains from the Barremian–Aptian transition (121 Ma, Hughes, 1994), which states the morphological age of angiosperms. Oldest pollen exhibiting subsets of definitive crown-angiosperm characters is known as far back as the Middle Triassic (237–247 Ma), but these are difficult to discriminate from pollen produced by stem-angiosperms or gymnosperms (Doyle and Hotton, 1991; Coiro et al., 2019). Slightly younger Aptian floral assemblages was followed by an explosive increase in diversity in the middle and Late Cretaceous (Lidgard and Crane, 1988; Friis et al., 2011). The fossil record provides fundamental evidence on the timescale and pattern of the origin and early evolution of angiosperms (Coiro et al., 2019). This is what some authors interpreted as an explosive radiation from a Cretaceous crown-ancestor (Benton et al., 2022). By contrast, there is a nearly universal molecular support for a pre-Cretaceous origin of crown angiosperms (Barba-Montoya et al., 2018).

Differences between molecular results and fossil record when estimating the age of crown angiosperms are not in conflict according to Sauquet et al. (2022), because both approaches rely ultimately on data from the fossil record since molecular dating studies are fossil-calibrated, albeit with different assumptions. Discrepancies between molecular clock and purely fossil-based interpretations of angiosperm diversification may be explained by two possible causes. One explanation suggests a long, cryptic evolutionary history of a non-ecologically significant group of early angiosperms (Doyle, 2012). They probably lived in environments in which fossilization was unlikely, resulting in an unrepresented fossil record. This idea is based on the early existence of triaperturate grain, known in the eudicots, which originated much later than the angiosperm-crown ancestor (Clarke et al., 2011). But this pollen character is also converging in Schizandraceae (Friis and Pedersen, 1996). It means that the record of the stem-group and original crown-group of angiosperms is still undiscovered, or at least unidentified. However, the synchronous diversification of fossil pollen, mesofossils and macrofossils through the Early Cretaceous would be difficult to explain if angiosperms had diversified cryptically for a significant time interval (Herendeen et al., 2017). It seems unlikely that crown groups emerge and keep an extensive period of hidden diversification. Instead, once they emerge, crown groups are probed to diversify rapidly and should quickly enter the fossil record (Budd and Mann, 2020). Thus, although early crown-group angiosperms may well have originated in the Jurassic (Coiro et al., 2019), the radiation of core angiosperms may have begun in the Cretaceous (Budd et al., 2021). An alternative explanation may be that molecular clock estimates are inaccurate with a trend to overestimation (Barba-Montoya et al., 2018). Indeed, molecular dating methods are not free from potential sources of bias (Beaulieu et al., 2015; Bromham et al., 2018; Budd et al., 2021; Sauquet et al., 2022). However, it is technically possible to reconciliate prior assumptions drawn from the fossil record with fossil-calibrated molecular dating approaches, since fossil-calibrated molecular dating analyses can be designed based on such prior assumptions of what crown angiosperm age is (Budd et al., 2021; Sauquet et al., 2022).

The discussion is taking part while putative crown-angiosperm fossils from the Jurassic of China (*e*.*g*., Fu et al., 2018) have been consistently rejected (Herendeen et al., 2017; Bateman, 2020; Sokoloff et al., 2020). All these supposed crown groups of angiosperms described from pre-Cretaceous time either represent other plant groups or lack features that might confidently assign them to the angiosperms (Coiro et al., 2019). It is therefore likely that some seed plants along the angiosperm stem-lineage from the Triassic and Jurassic possessed some, but not all, of the features of crown group angiosperms. So far, however, the discrepancy remains, due to the lack of undisputed pre-Cretaceous crown-angiosperm fossils (Herendeen et al., 2017).

Despite the uncertainty in the timing of the origin of crown-angiosperms, it seems clear that the diversification of the major lineages of angiosperms occurred during the Late Cretaceous (Magallón et al., 2015; Barba-Montoya et al., 2018; Li et al., 2019; Budd et al., 2021). From that time, the fossil record reflects flowering plants to have risen to ecological dominance in terrestrial communities up to the Paleogene (Silvestro et al., 2015; Ramírez-Barahona et al., 2020), along with the origin of hyperdiverse biomes such as tropical rainforests (Carvalho et al., 2021). This time does appear to coincide with a peak in the diversity of insects, including herbivores and pollinators and their predators (Claphman et al., 2016; Schachat et al., 2019), corroborating the hypothesis of the ATR — an explosive boost to terrestrial diversity occurred from 100–50 Ma, the Late Cretaceous and early Palaeogene (Benton et al., 2022).

### Evolution of pollinators

Assuming that a pollinator is an organism that moves pollen from the male anther to the female stigma in flowers is simplistic. This statement faces two main problems. On one hand, insect pollination (entomophily) is a process that occurs especially but not exclusively on flowering plants. While around 85% of flowering plants are biotically pollinated (Ollerton et al., 2011), around 40% of gymnosperms (including Cycadales and Gnetales) are also pollinated by insects (Ickert-Bond and Renner, 2016; Toon et al., 2020). However, pollination requirements of most gymnosperms and most wild plants are surprisingly unknown. On the other hand, if the definition of a pollinator implies an active process, it would be difficult to find evidence of such a behaviour from the fossil record, and when and how the pollination process evolved. The characters that a fossil must meet to be considered a pollinator is still an open discussion (Peris et al., 2020; Peña-Kairath et al., in prep.).

Organisms involved in the pollination process include iconic groups of Recent pollinators, namely bees (Hymenoptera), long-proboscis butterflies (Lepidoptera), or many different groups of beetles and flies (Coleoptera and Diptera, respectively). However, although less significant in present times, some families in Thysanoptera, Hemiptera, Neuroptera, Orthoptera and Blattodea have been proved to be pollinators of some plant groups (Ollerton et al., 2017; Peña-Kairath et al., in prep.) (Figure 4). Within the extant insect orders, different extinct groups in Neuroptera, Mecoptera and Diptera display long mouthparts and rostra adapted to feeding on pollination drops of extinct gymnosperms and floral nectar (Khramov et al., 2020, 2022). It is interesting to note that all ancient representatives with long mouthparts are currently extinct in Mecoptera and Neuroptera, and that Mecoptera, unlike Neuroptera, do not include specialized-pollen feeders today. This indicates that any relationship with pollination in Mecoptera corresponds to an ancestral behaviour. Finally, two fossil groups of insects have also been cited as presumable pollinators in deep times (Peña-Kairath et al., in prep.): Permopsocida, an insect order known from the early Permian but extinct during the mid-Cretaceous, for which angiosperm pollen has been found in the intestines and in contact with the abdomen of a taxon (Huang et al., 2016); and Alienoptera, an extinct group with described species since the Early Cretaceous to the Eocene, for which a Cretaceous nymph has been described in contact with gymnosperm pollen clumps (Luo et al., 2022). Fossils from the extant order Grylloblattodea and the extinct orders Miomoptera and Hypoperlida have been described form early Permian sediments, more than 270 Ma, with pollen grains in their guts (references in Peña-Kairath et al., in prep.), which suggests these organisms where likely consuming pollen at that time. However, their pollination habit in these last cases is more dubious (Peña-Kairath et al., in prep.).

**Figure 4.**
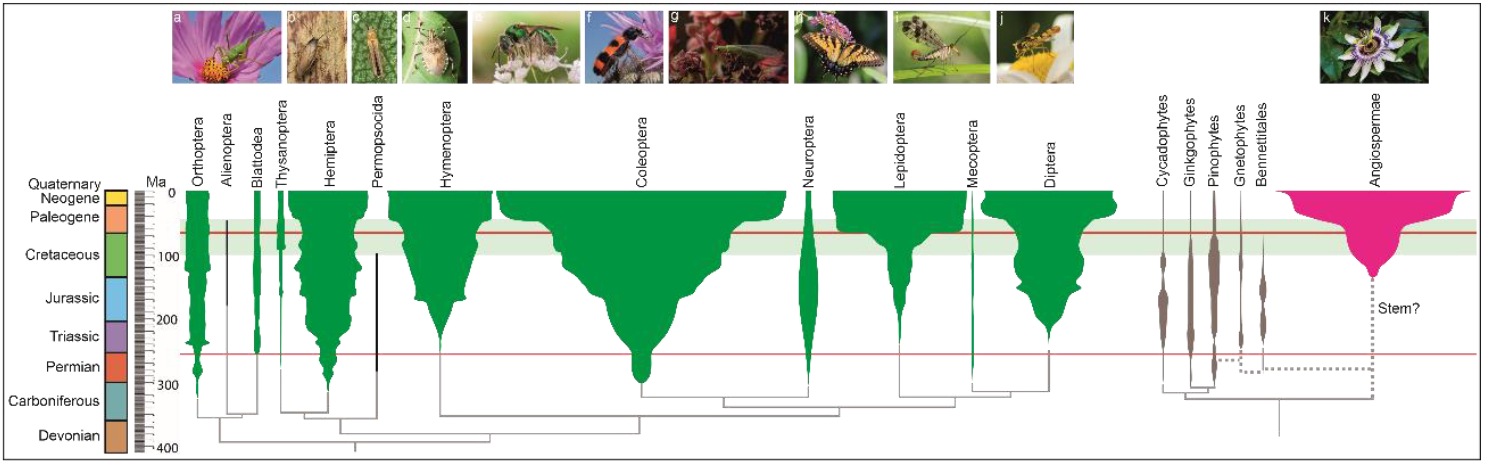
Evolution of the insect orders with extant and/or extinct pollinator representatives, gymnosperms and angiosperms (flowering plants). Figure based on the “Tree of life” from the Museum of Natural History at the Oxford University, courtesy from Dr. Ricardo Pérez de la Fuente. Each of the lineages shown in the tree is scaled to the described number of extant species. Species’ recounts taken from the CoL (https://www.catalogueoflife.org/). Recent groups spindles of insects extracted from Labandeira and Eble (2000) and of plants from Benton et al. (2022). As Nicholson et al. (2015) showed, since 2000 the curve of insect diversity -based on families-through time has not substantially changed. Smaller lineages are namely represented as a line. Phylogenetic relationships of insects extracted from Misof et al. (2014), Beutel et al. (2017), and Giribet and Edgecombe (2019). Phylogenetic relationships of plants extracted from Novikov and Barabasz-Krasny (2015). Divergence times obtained using the TimeTree website http://www.timetree.org/ except for the fossil insect orders Permopsocida (Huang et al., 2016) and Alienoptera (Luo et al., 2022). The origin of the Gnetophytes and the extinct Bennetitalean plants is controversial. The TimeTree website collates multiples papers dealing with divergence times and obtains an average. Red lines represent significant mass extinctions. Green zone represents the Angiosperm Terrestrial Revolution after Benton et al. (2022). Royalty-free images of insects and angiosperm on the top of the figure obtained from the website www.freepick.es: a. Orthoptera; b. Blattodea; c. Thysanoptera; d. Hemiptera; e. Hymenoptera; f. Coleoptera; g. Neroptera; h. Lepidoptera; i. Mecoptera; j. Diptera; k. Angiospermae.

The fossil record together with phylogenetic studies show a scenario in which all the extant and extinct orders of insects that include pollinators evolved well before the Early Cretaceous (Figure 4). They show an origin that largely predates the diversification of crown angiosperms, except for lineages of some flies, bees and long-proboscid butterflies (Supplementary Information).

### Angiosperms evolved in a time of insect diversity peak

A growing number of fossil insect taxa have been described pollinating different groups of plants in the fossil record. All these fossil pollinators were described exclusively associated with gymnosperms until the beginning of the Late Cretaceous, when a combination of both gymnosperm and angiosperm hosts are found associated with the described fossils, but only evidence of angiosperm host remain in more recent cases (Peris et al., 2020; Peña-Kairath et al., in prep.) (Figure 1). Entomophilous pollination is found to be common during the Cretaceous, which is consistent with the diversification of the major groups of angiosperms around 100 Ma (Barba-Montoya et al., 2018; Budd et al., 2021) and their rise to ecological dominance in terrestrial communities in the late Cenozoic (Condamine et al. 2020; Ramírez-Barahona et al., 2020).

Highly specialized pollination is evidenced in some groups of insects by the development of suctorial feeding structures such as long-proboscid nectar-feeding mouthparts (Khramov et al., 2020), described since the early Permian (Khramov et al., 2022). Such mouthparts are thought to be used for feeding on pollination drops released on the micropyles of the ovules concealed inside the gymnosperm strobili (Labandeira et al., 2007). It is not surprising that lineages with specific adaptations were extinct together with their gymnosperm host when angiosperms turnover ancient ecosystems, such as the long-proboscid Mecoptera, Neuroptera, and the zhangsolvid Diptera (Figure 1). Plant diversity declined by 45% at the K/Pg boundary (Silvestro et al., 2015; Condamine et al. 2020; Carvalho et al., 2021) probably exacerbated by pollinator specialization. By contrast, other groups represent generalist feeding habit, such as mandibulate beetles, that successfully transitioned from a gymnosperm- to angiosperm-dominated flora (Peris et al., 2017, 2020). With the beginning of the Cenozoic the long-proboscid Mecoptera and Neuroptera were supplanted by Hymenoptera and Lepidoptera in the nectar-feeding niche (Khramov et al., 2020), but already on angiosperms. All these cases correspond to different pollination modes covered in Peris et al. (2017), showing different groups of insects under distinctive patterns of extinction, survival, and origination following the gymnosperm-angiosperm transition (Figure 1). Wind pollination probably evolved in angiosperms from insect pollination in response to pollinator limitation and changes in the abiotic environment (Culley et al., 2002). Accordingly, insect pollination is still considered as the initial pollination mode for angiosperms (Hu et al., 2008).

Growing evidence from molecular dated phylogenetics, fossil record of pollinator insects, paleontological data on morphological features of plants and modelling of diversification dynamics support the hypothesis that angiosperms evolved in a time of insect diversity peak during the Early Cretaceous (Claphman et al., 2016; Schachat et al., 2019) (Figure 2). The essential trophic machinery of insects was in place nearly 100 million years before angiosperms appeared in the fossil record (Labandeira and Sepkoski, 1993). This high diversity represents a true burst of origination (Schachat et al., 2019), including insect lineages with highly adapted, pollination modes on gymnosperms (Peris et al., 2020; Khramov et al., 2020). That would mean that gymnosperm pollinators were available for angiosperms when they evolved, predating the first flowering plants (Labandeira, 2007; Peris et al., 2017, 2020; Khramov et al., 2020). However, assuming that the fossil record might be biased and incomplete, flowers might have arisen earlier according to molecular phylogenetics of plants. In addition, pollen and pollination-fluid-feeding have evolved independently across more than a dozen of insect orders, and in hundreds of families (Wardhaugh, 2015). Such an evolutionary convergence supports the hypothesis that it is relatively easy for pollinivory to evolve from feeding on the spores of fungi, ferns and other spore plants (Labandeira, 2000). The pre-Cretaceous diversity of insects that developed with non-angiosperm plants might have constituted a supertramp by which adaptations arose when angiosperms diversified and replaced gymnosperms by the end of the Cretaceous (Condamine et al. 2020).

### Role of the Angiosperm Revolution on the insect diversification

The apparent temporal gap between insect pollinators and crown angiosperms seems to question the reciprocal diversification model, or coevolution between insects and flowering plants (Asar et al., 2022). Relying on qualitative comparisons has limits, and there are little quantitative tests to evaluate whether coevolution has spurred diversification between plants and insects. Here we retrieved the family-level fossil dataset of Condamine et al. (2016) to estimate whether origination and/or extinction rates varied through time and whether they could be correlated to the angiosperm radiation. We relied on the Bayesian framework implemented in PyRate (Silvestro et al., 2014, 2019) to simultaneously estimate correlations between diversification dynamics and multiple environmental variables (Lehtonen et al., 2017). We focus on the role of angiosperms but are aware that several drivers can have complementary impacted the diversification of insects (see *Methods* for details). We thus incorporated six variables: the number of insect family through time (diversity dependence), the relative diversity of angiosperms, gymnosperms and spore plants (Silvestro et al., 2015), the continental fragmentation (Zaffos et al., 2017), and the change in temperature (Prokoph et al., 2008).

First, we estimated origination and extinction rates across the whole evolutionary history of insects (*i*.*e*., mid-Carboniferous to present), for which angiosperms were absent until the Early Cretaceous. Our modelling results shows that, among the six tested variables, angiosperms promoted a faster diversification of insects once they coevolved since the Cretaceous (Table 1a). This remarkable link between insect origination with angiosperm radiation was most probably the cause that drove the radiation of different groups of herbivores such as beetles (McKenna et al., 2009; Ahrens et al., 2014; Doorenweerd et al., 2017) and pollinators such as bees and long-proboscid butterflies since the mid-Cretaceous onwards (Cardinal and Danforth, 2013; Chazot et al., 2019). In addition, we found that the rise of flowering plants generally not only drove the origination of insect families, but they also buffer them against extinction (Table 1a). We then performed the same diversification analyses but with rates only estimated for the time interval covering the ATR (*sensu* Benton et al., 2022, from 100 to 50 Ma) and another set of analyses with rates only estimated in the aftermath of the ATR (from 50 Ma to present). The ATR-centred analyses indicated that the rise of angiosperms strongly decreased insect extinction rates during the ATR but did not affect insect origination rates (Table 1b, Figure 3). The post-ATR analyses indicated that the rise of angiosperms strongly increased insect origination rates and decreased insect extinction rates, but lesser than during the ATR (Table 1c, Figure 3). Therefore, angiosperms had a dual role that has changed through time with an attenuation of the extinction in the Cretaceous and beginning of the Cenozoic and a driver of origination in the Cenozoic, from 50 Ma onwards.

It is also important to highlight, nevertheless, that the rise of angiosperms is not the only driver with significant identified effect on insect evolution (Table 1, Figure 3). The analyses also indicate that increasing insect diversity has slowed down origination rates in insects, which is line with the role of diversity dependence. Spore plants are also found as a strong positive correlation with both origination and extinction rates of insects, suggesting that higher spore plant diversity spurred insect turnover; a result that would deserve further studies given the important effect in our analyses (Table 1). Gymnosperms are never recovered as a significant driver in our analyses. We also unveil that global temperature correlates negatively with origination and positively with extinction, such that warmer climate led to lower origination and higher extinction in insects, which partly corroborates previous results for invertebrates (Mayhew et al., 2008). Different alternative hypotheses are also cited in the literature (*eg*. in Labandeira, 2014) but not covered in this analysis, which is focused eminently on the angiosperm-insect pollinators coevolution.

### Triggers for the angiosperm success

The vast current diversity of angiosperms has been ascribed to different innovations in their reproductive, vegetative and genomic biology, which presumably played a central role in their diversification and rise to ecological dominance (Vamosi and Vamosi, 2011; Sauquet and Magallón, 2018; Vamosi et al., 2018; Benton et al., 2022). These attributes were considered to provide reproductive superiority over non-angiosperm seed plants, favouring their evolution after their origin (Soltis et al., 2019). Although diversification rate shifts are caused by changes in both speciation and extinction rates, it is not trivial to find the causes of rate shifts in such a biological radiation.

Many of the morphological features thought to represent key innovations in angiosperms may not be the only, or even the primary, cause of diversification rate shifts. The trait and ecological shifts driving higher rates of diversification have not generally been the synapomorphies used by systematists to define the major clades of angiosperms, such as flowers, double fertilization, endosperm development, and seeds encased in fruits (Vamosi et al., 2018). Instead, intrinsic key innovations, extrinsic factors such as geography and environment, or trait– environment combinations are important drivers of diversification rates (Vamosi and Vamosi, 2011; Chaboureau et al.; 2014; Bouchenak-Khelladi et al., 2015; Lagomarsino et al., 2016; O’Meara et al., 2016; reviewed in Vamosi et al., 2018; Simonin and Roddy, 2018; Roddy et al., 2020). Intrinsic key innovations include the whole-genome duplications event, named polyploidy, shared by all living angiosperms; the ability to reduce genome size, that enabled the reduction of the cell sizes; the herbaceousness, that is short generation time; maximizing photosynthetic efficiency by the increase of venation and stomatal density into their leaves since ∼100 Ma (see references from above). Extrinsic factors can include patchy habitats, with habitat generalization leading to decreased extinction, open habitats favouring diversification of lineages adapted to such conditions, and island-like regions (including mountainous regions) increasing rates of allopatric speciation.

Close associations of flowering plants with pollinator insects are particularly supposed to have played an early role in angiosperm diversification (Crepet 1984; Grimaldi 1999; Hu et al. 2008; Van der Niet et al., 2014). This co-diversification seemed to be the result from a pollinator transition of generalist pollen-feeding insects from gymnosperms to angiosperms (Labandeira et al., 2007; Peris et al., 2017, 2020). However, this hypothesis was called into question because the advantages offered by the gymnosperm pollinators did not prevent them from a decline (Crisp and Cook, 2011; Condamine et al., 2020; Gorelick, 2001; Khramov et al., 2020) and no plants other than angiosperms are diverse nowadays. Thus, it would be necessary to analyse the drivers of diversification in angiosperms (O’Meara et al., 2016) using a similar approach that we applied for insects here. While insect diversification was driven faster thanks to the coadaptation to angiosperms, our analysis shows that the gymnosperms are not recovered as a significant factor for the insect diversification (Table 1, Figure 3).

The rise of flowering plants through the Cretaceous, twice as productive as gymnosperms (Boyce and Lee, 2017), was followed by new systems of chemical defense, tolerance to climatic stress, and the (genetic) ability of certain angiosperm lineages to repeatedly evolve adaptive traits (Onstein, 2020). Those facts, together with climate changes, breakup of Pangea, increase of humid conditions during the Late Cretaceous (Chaboureau et al., 2014) and the global cooling at the end of the Paleogene, have been linked to the decline in conifer diversity from the Cretaceous (the last 110 Ma) and extended through the Cenozoic (Crisp and Cook, 2011; Condamine et al., 2020). The rise of angiosperms induced an active displacement by outcompeting conifers (Condamine et al., 2020), and caused conifers to have high extinction rates ever since (Crisp and Cook, 2011). Faced with this situation, gymnosperm pollinators likely had little options, adapt or go extinct depending on how specialized they were. The once diverse Cheirolepidiaceae and Bennettitales went extinct around the K/Pg boundary, as did some highly specialized long-proboscid flies, scorpionflies and lacewings related to these plants (Rothwell et al., 2009; Khramov et al., 2020).

Angiosperms did not achieve ecological dominance in a single step (Davies et al., 2004; Magallón and Castillo, 2009; Onstein, 2020). They were still low-biomass components of most Cretaceous floras (Carvalho et al., 2021). Rather, the appearance of various traits, the ability of certain angiosperm lineages to repeatedly evolve them, clade-specific radiations (Sauquet et al., 2017; Onstein, 2020; Ramírez-Barahona et al., 2020), and their link with insect diversification likely spurred diversity in different groups following the K/Pg event (Carvalho et al., 2021). It was in the aftermath of the K/Pg event that the diversification of angiosperms and of insects had explosive impacts on each other through their species interactions (Asar et al., 2022), and only then angiosperms achieved ecological dominance (Carvalho et al., 2021). The diversity and dominance of crown-group angiosperm families and genera continued to rise in the Cenozoic, accelerating insect-angiosperm co-diversification (Allio et al., 2021), evolving multiple lineages of pollinating insects (Wiegmann et al., 2011; Sann et al., 2018; Chazot et al., 2019; Song et al., 2020; Cai et al., 2022), driving to the ‘modernization’ of Earth ecosystems.

### Future considerations

We still know little about the origins of entomophily and how it evolved, but theories regarding pollination-plant coevolution always predict an increased probability of radiation of both plants and the pollinating animals because of the mutualistic nature of the interaction (Gorelick, 2001). However, it does not always hold. Coevolutionary processes should not be considered the only major drivers of diversification in plants and insects (Suchan and Alvarez, 2015), which is confirmed by our analyses indicating that global temperature, spore plants, and diversity dependence are additional drivers to study for explaining insect diversity. For instance, warmer global temperature drove higher origination and extinction in insects (Currano et al., 2016). Insect pollination is not a guarantee of higher success. For example, cycads are insect pollinated and have never been a diverse lineage (Condamine et al., 2015; Toon et al., 2020). By getting more specialized, it also increases the pollination efficiency, providing a mechanism to explain the boost in speciation rates (Citerne et al., 2010). However, specialization also increases the risk of extinction rate with environmental upheavals. Under fluctuating conditions, plants that are pollinated by specific animals will be more adversely affected than plants that are generally pollinated by several species. Most of current biotically pollinated plants and their pollinating animals are generalists (Waser et al., 1996), although insect pollinators that feed from and pollinate a single plant species do exist (Gorelick, 2001; Toon et al., 2020). The angiosperm extinction rates decreased after the K/Pg boundary in parallel to an increased speciation (Silvestro et al., 2015), while the contrary is observed with the conifer diversity (Condamine et al., 2020).

## Conclusions

The origin of angiosperms and their coevolution with potential pollinators remains enigmatic, but significant progresses have been made in the last decade with fossil-based and phylogenetic studies. Early diversification of angiosperms and their insect pollinators were largely decoupled (van der Kooi and Ollerton, 2020; Asar et al., 2022). Pollinator insect lineages predated flowers (Labandeira et al., 2007; Peris et al., 2020; Asar et al., 2022) (Figure 4). We also know that insect family-richness peaked 125 Ma (Claphman et al., 2016; Schachat et al., 2019) (Figure 2), which coincides with numerous pollinator lineages that were adapted to pollinate gymnosperms (Labandeira et al., 2007; Peris et al., 2017; Peris et al., 2020). By contrast, most angiosperm families (58–80%) originated between ∼100 and 90 Ma, during the warmest phases of the Cretaceous (Ramírez-Barahona et al., 2020). This is also the exact time when first angiosperm pollinators are found in the fossil record (Peris et al., 2020; Peña-Kairath et al., in prep.).

Despite their time of origin, the rise to ecological dominance of modern-day angiosperms was geographically heterogeneous and took place after a prolonged period, lasting until the Palaeocene, concomitant with the onset of crown diversification in most families (Crisp and Cook, 2011; Ramírez-Barahona et al., 2020), and pushing different gymnosperm lineages to decline (Condamine et al. 2020; Mazet et al., 2022). We have found that the flowering plant evolution promoted a faster diversification of insects, not only driving the origination of insect families but also buffering them against extinction since the mid-Cretaceous (Table 1, Figures 2–3).

Reducing the idea of angiosperm-pollinator co-evolution to a single period under the analysis of being cause or consequence of each other (*e*.*g*., Khramov et al., 2020) may be an excessive simplification. On one hand, there was a significant pool of gymnosperm pollinators that might have been available for angiosperms since their beginning (Labandeira et al., 2007; Peris et al., 2017; Peris et al., 2020). On the other hand, we found a link between the diversification of angiosperms and insects, including the evolution of Late Cretaceous pollinator lineages such as bees and butterflies (Sann et al., 2018; Chazot et al., 2019). Pollination is a very complex system of mutualistic relationships that is necessary to be analysed in time, space and morphology case by case (*eg*., Lavaut et al., 2022). Focusing on a single driver of increased diversification seems to be a reductionist approach that will lead to incomplete explanations (O’Meara et al. 2016; Sauquet and Magallón, 2018). More likely, trait–environment combinations may spur diversification affecting speciation and/or extinction rates (Bouchenak-Khelladi et al., 2015; Lagomarsino et al., 2016; O’Meara et al., 2016; Freyman and Höhna, 2019).

## Methods

### Fossil data and multivariate birth-death analyses

We retrieved the times of origination and times of extinction for 1,527 families of which 671 extant and 856 extinct families, which have been estimated from 38,000+ fossil occurrences at the family level (Condamine et al., 2016). To examine variations of insect family diversity through time, we reconstructed the lineages-through-time using PyRate 3 (Silvestro et al., 2019) and the input file of origination and extinction times of all insect families (*-ltt 1* option).

PyRate has developed and implemented this birth-death model to test for a correlation between speciation and extinction rates and changes in environmental variables (Lehtonen et al., 2017). We used the Multivariate Birth-Death model (MBD) to assess to what extent biotic and abiotic factors can explain temporal variation in speciation and extinction rates. Under the MBD model, speciation and extinction rates can change through time, but equally across all lineages, through correlations with multiple time-continuous variables, and the strengths and signs (positive or negative) of the correlations are jointly estimated for each variable (Lehtonen et al., 2017). The MBD model includes temporal fluctuations of environmental variables, so that the speciation and extinction rates can depend on the variations of each factor. The correlation parameters can take negative values indicating negative correlation, or positive values for positive correlations. When their value is estimated to be approximately zero, no correlation is estimated. A Markov chain Monte Carlo (MCMC) algorithm jointly estimates the baseline speciation (λ0) and extinction (μ0) rates and all correlation parameters (*G*λ and *G*μ) using a horseshoe prior to control for over-parameterization and for the potential effects of multiple testing. The horseshoe prior provides an efficient approach to distinguishing correlation parameters that should be treated as noise (and therefore shrunk around 0) from those that are significantly different from 0 and represent true signal. We ran the MBD model using 50 million MCMC iterations and sampling every 50,000 to approximate the posterior distribution of all parameters (λ0,μ 0, ten *G*λ, ten *G*μ, and the shrinkage weights of each correlation parameter, ωG). We summarized the results of the MBD analyses by calculating the posterior mean and 95% credibility interval of all correlation parameters and the mean of the respective shrinkage weights (across ten replicates), as well as the mean and 95% credibility interval of the baseline speciation and extinction rates.

The MBD model assumes that diversification rates vary continuously through time with a given variable. However, it is possible that rates vary positively during a given time interval and then negatively in another time interval (Neubauer et al., 2022). In other words, drivers of diversification can vary over time. We thus tested whether the impact of angiosperm diversity was similar over time, during the ATR (100-50 Ma, *sensu* Benton et al. 2022) and post-ATR (50-0 Ma). The MBD analyses were thus performed with time constraints to estimate rates within this time interval by setting up the *-maxT 100 minT 50* or *-maxT 50 minT 0* option to represent the ATR timeframe and the post-ATR period, respectively.

### Paleoenvironmental variables

To identify putative mechanisms of insect diversification, we examined the correlation between a series of past environmental variables and origination/extinction rates over their entire history. We focused on the role of four paleoenvironmental variables, also called proxies, which have been linked to biodiversity change. These proxies were classified as either abiotic or biotic controls as follows: *(i) Abiotic controls*. Climate change (variations from warming to cooling periods) is one of the most probable drivers of diversification changes throughout the history of life (Erwin, 2009; Condamine et al., 2019). Major trends in global climate change through time are typically estimated from relative proportions of different oxygen isotopes (δ^18^O) in samples of benthic foraminifer shells. We merged δ^18^O global temperature data from different sources (Westerhold et al., 2020 for the Cenozoic; Veizer and Prokoph, 2015 for the rest of the Phanerozoic) to provide δ^18^O data spanning the full time-range over which insect families originated. Second, global continental fragmentation, as approximated by plate tectonic change over time, has also been proposed as a driver of diversity dynamics (Zaffos et al., 2017). We retrieved the index of continental fragmentation developed by Zaffos et al. (2017) using paleogeographic reconstructions for 1-million-year time intervals. This index approaches 1 when all plates are not connected (complete plate fragmentation) and approaches 0 when there is maximum aggregation. *(ii) Biotic controls*. Ecological interactions with rapidly expanding clades are increasingly recognized as important macroevolutionary drivers (Condamine et al., 2020). Insects experienced drastic floristic changes throughout their evolutionary history with the origin and rapid radiation of angiosperms at the expense of a decline in diversity of gymnosperms and ferns. The rise and dominance of angiosperms may have contributed to altering the dietary regimes of herbivorous insects, which could in turn have affected insects that depend on herbivores by a cascading effect. We thus compiled the relative diversity trajectories of angiosperms, gymnosperms, and spore plants (mostly composed of ferns) based on previous estimates of plant diversity (Silvestro et al., 2015). *(iii) Insect diversity*. Biotic interactions within insects could also have influenced their diversification. For instance, we could draw hypotheses of diversity dependence such that insects could either impact or be impacted by their own diversity. In other words, the change in their diversity can affect their diversification as proposed in diversity-dependent hypotheses. We thus included the paleodiversity of all insect families to account for diversity dependence within insects as an independent variable.

### Figures

Figures were created using CorelDRAW Graphics Suite software, version 19.0. (www.coreldraw.com). Figure 2 was designed based on the results obtained after the data analysis with the software R. Figure 4 was designed based on the “Tree of life” from the Museum of Natural History at the Oxford University, courtesy from Dr. Ricardo Pérez de la Fuente.

## Acknowledgements

DP thanks the Ministry of Economy and Competitiveness of Spain (project “CRE”, Spanish AEI/FEDER, UE CGL2017-84419) for financial support. This is contribution no. 4 of the postdoctoral fellowships programme Beatriu de Pinós project 2020 BP 00015, *The flowering plant success – Influence of beetles*, funded to DP by the Secretary of Universities and Research (Government of Catalonia) and by the Horizon 2020 programme of research and innovation of the European Union under the Marie-Curie grant agreement no. 801370. FLC is supported by the European Research Council (ERC) under the European Union’s Horizon 2020 research and innovation programme (project GAIA, agreement no. 851188).

## Data and materials availability

All data are available in the main text or the supplementary information.

## The following Supporting Information is available for this article

Supplementary Information – Origination of the insect pollinator lineages

## References

1. Ahrens, D., Schwarzer, J., Vogler, A.P. 2014 The evolution of scarab beetles tracks the sequential rise of angiosperms and mammals. Proceedings of the Royal Society B: Biological Sciences, 281, 20141470.

2. Allio, R., Nabholz, B., Wanke, S., Chomicki, G., Pérez-Escobar, O.A., Cotton, A.M., CLamens, A.L., Kergoat, G.J., Sperling, F.A.H., Condamine, F.L. 2021. Genome-wide macroevolutionary signatures of key innovations in butterflies colonizing new host plants. Nature Communications, 12, 354.

3. Asar, Y., Ho, S.Y.W., Sauquet, H. 2022. Early diversifications of angiosperms and their insect pollinators: were they unlinked? Trends in Plant Science, 27, 858–869.

4. Barba-Montoya, J., dos Reis, M., Schneider, H., Donoghue, P., Yang, Z. 2018. Constraining uncertainty in the timescale of angiosperm evolution and the veracity of a Cretaceous Terrestrial Revolution. New Phytologist, 218, 819–834.

5. Bateman, R.M. 2020. Hunting the Snark: the flawed search for mythical Jurassic angiosperms. Journal of Experimental Botany, 71, 22–35.

6. Benton, M.J., Wilf, P., Sauquet, H. 2022. The Angiosperm Terrestrial Revolution and the origins of modern biodiversity. New Phytologist, 233, 2017–2035.

7. Bouchenak-Khelladi, Y., Onstein, R.E., Xing, Y., Schwery, O., Linder, H.P. 2015. On the complexity of triggering evolutionary radiations. New Phytologist, 207, 313–26.

8. Boyce, C.K., Lee, J.E. 2010. An exceptional role for flowering plant physiology in the expansion of tropical rainforests and biodiversity. Proceedings of the Royal Society B: Biological Sciences, 277, 3437–3443.

9. Bromham, L., Duchêne, S., Hua, X., Ritchie, A.M., Duchêne, D.A., Ho, S.Y. 2018. Bayesian molecular dating: opening up the black. Biological Reviews, 93, 1165–1191.

10. Budd, G.E., Mann, R.P. 2020. The dynamics of stem and crown groups. Science Advances, 6, eaaz1626.

11. Budd, G.E., Mann, R.P., Doyle, J.A., Coiro, M., Hilton, J. 2021. Fossil data do not support a long pre-Cretaceous history of flowering plants. bioRxiv, 2021.02.16.431478.

12. Buggs, R.J.A. 2021. The origin of Darwin’s “abominable mystery.” American Journal of Botany, 108, 22–36.

13. Cai, C., Escalona, H.E., Li, L., Yin, Z., Huang, D., Engel, M.S. 2018. Beetle pollination of cycads in the Mesozoic. Current Biology, 28, 2806–2812.

14. Cai, C., Tihelka, E., Giacomelli, M., Lawrence, J.F., Slipinski, A., Kundrata, R., Yamamoto, S., Thayer, M.K., Newton, A.F., Leschen, R.A.B., Gimmel, M.L., Engel, M.S., Boucharcd, P., Huang, D., Pisani, D., Donoghue, P.C.J. 2022. Integrated phylogenomics and fossil data illuminate the evolution of beetles. Royal Society Open Science, 9, 211771.

15. Christenhusz, M.J.M., Byng, J.W. 2016. The number of known plants species in the world and its annual increase. Phytotaxa, 261, 201–217.

16. Clapham, M.E., Karr, J.A., Nicholson, D.B., Ross, A.J., Mayhew, P.J. 2016. Ancient origin of high taxonomic richness among insects. Proceedings of the Royal Society B: Biological Sciences, 283, 20152476.

17. Clarke, J.T., Warnock, R.C.M., Donoghue, P.C.J. 2011. Establishing a time-scale for plant evolution. New Phytologist, 192, 266–301.

18. Cardinal, S., Danforth, B.N. 2013. Bees diversified in the age of eudicots. Proceedings of the Royal Society B: Biological Sciences, 280, 20122686.

19. Carvalho, M.R., Jaramillo, C., de la Parra, F.,Caballero-Rodríguez, D., Herrera, F., Wing, S., Turner, B., D’Apolito, C., Romera-Báez, M., Narváez, P., Martínez, C., Gutiérrez, M., Labandeira, C.C., Bayona, G., Rueda, M., Paez-Reyes, M., Álvaro-Duque, D., Crowley, J.L., Santos, C., Silvestro, D. 2021. Extinction at the end-Cretaceous and the origin of modern Neotropical rainforests. Science, 372, 63–68.

20. Citerne, H.L., Jabbour, F., Nadot, S., Damerval, C. 2010. The evolution of floral symmetry. Advances in Botanical Research, 54, 85–137.

21. Claphman, M.E., Karr, J.A., Nicholson, D.B., Ross, A.J., Mayhew, P.J. 2016. Ancient origin of high taxonomic richness among insects. Proceedings of the Royal Society B: Biological Sciences, 283, 20152476.

22. Crepet, W.L. 1984. Advanced (constant) insect pollination mechanisms: pattern of evolution and implications vis-á-vis angiosperm diversity. Annals of the Missouri Botanical Garden, 71, 607–630.

23. Coiro, M., Doyle, J.A., Hilton, J. 2019. How deep is the conflict between molecular and fossil evidence on the age of angiosperms? New Phytologist, 223, 83–99.

24. Condamine, F.L., Nagalingum, N.S., Marshall, C.R., Morlon, H. 2015. Origin and diversification of living cycads: a cautionary tale on the impact of the branching process prior in Bayesian molecular dating. BMC Evolutionary Biology, 15, 1–18.

25. Condamine, F.L., Clapham, M.E., Kergoat, G.J. 2016. Global patterns of insect diversification: towards a reconciliation of fossil and molecular evidence? Scientific Reports, 6, 19208.

26. Condamine, F.L., Silvestro, D., Koppelhus, E.B., Antonelli, A. 2020. The rise of angiosperms pushed conifers to decline during global cooling. Proceedings of the National Academy of Sciences of the United States of America, 117, 28867–28875.

27. Crisp, M.D., Cook, L.G. 2011. Cenozoic extinctions account for the low diversity of extant gymnosperms compared with angiosperms. New Phytologist, 192, 997–1009.

28. Culley, T.M., Weller, S.G., Sakai, A.K. 2002, The evolution of wind pollination in angiosperms. Trends in Ecology & Evolution, 17, 361–369.

29. Currano, E.D., Laker, R., Flynn, A.G., Fogt, K.K., Stradtman, H., Wing, S.L. 2016. Consequences of elevated temperature and pCO2 on insect folivory at the ecosystem level: perspectives from the fossil record. Ecology and Evolution, 6, 4318–4331.

30. Davies, T.J., Barraclough, T.G., Chase, M.W., Soltis, P.S., Soltis, D.E., Savolainen, V. 2004. Darwin’s abominable mystery: insights from a supertree of the angiosperms. Proceedings of the National Academy of Sciences of the United States of America, 101, 1904–1909.

31. Doorenweerd, C., Van Nieukerken, E.J., Hoare, R.J. 2017. Phylogeny, classification and divergence times of pygmy leaf-mining moths (Lepidoptera: Nepticulidae): the earliest lepidopteran radiation on Angiosperms? Systematic Entomology, 42, 267–287.

32. Doyle, J.A. 2012. Molecular and fossil evidence on the origin of angiosperms. Annual Review of Earth and Planetary Sciences, 40, 301–326.

33. Doyle, J.A., Hotton, C.L. 1991. Diversification of early angiosperm pollen in a cladistic context. In: Blackmore S, Barnes S, (Eds.) Pollen and spores: patterns of diversification. Oxford, UK: Clarendon Press, 169–195.

34. Erwin, D.H. 2009. Climate as a driver of evolutionary change. Current Biology, 19, 575–583.

35. Friis, E.M., Pedersen, K.R. 1996. Eucommiitheca, a new pollen organ with Eucommiidites pollen from the Early Cretaceous of Portugal. Grana, 35, 104–112.

36. Friis, E.M., Crane, P.R., Pedersen, K.R. 2011. Early Flowers and Angiosperm Evolution Cambridge University Press.

37. Fu, Q., Diez, J.B., Pole, M., García-Ávila, M., Liu, Z., Chu, H., Hou, Y., Yin, P., Zhang, G., Du, K., Wang, X. 2018. An unexpected noncarpellate epigynous flower from the Jurassic of China. eLife, 7, 1–24.

38. Giribet, G., Edgecombe, G.D. 2019. The phylogeny and evolutionary history of arthropods. Current Biology, 29, 592–602.

39. Grimaldi, D. 1999. The co-radiations of pollinating insects and angiosperms in the Cretaceous. Annals of the Missouri Botanical Garden, 86, 373–406.

40. Gorelick, R. 2001. Did insect pollination cause increased seed plant diversity? Biological Journal of the Linnean Society, 74, 407–427.

41. Herendeen, P.S., Friis, E.M., Pedersen, K.R., Crane, P.R. 2017. Palaeobotanical redux: revisiting the age of the angiosperms. Nature Plants, 3, 17015.

42. Hu, S., Dilcher, D.L., Jarzen, D.M., Taylor, D.W. 2008. Early steps of angiosperm– pollinator coevolution. Proceedings of the National Academy of Sciences of the United States of America, 105, 240–245.

43. Huang, D.-Y., Bechly, G., Nel, P., Engel, M.S., Prokop, J., Azar, D., Cai, C.-Y., van de Kamp, T., Staniczek, A.H., Garrouste, R., Krogman, L., dos Santos Rolo, T., Baumabach, T., Ohlhoff, R., Shmakov, A.S., Bourgoin, T. Nel, A. 2016. New fossil insect order Permopsocida elucidates major radiation and evolution of suction feeding in hemimetabolous insects (Hexapoda: Acercaria). Scientific Reports, 6, 23004.

44. Hughes, N.F. 1994. The Enigma of Angiosperm Origins (Cambridge University Press).

45. Ickert-Bond, S.M., Renner, S.S. 2016. The Gnetales: recent insights on their morphology, reproductive biology, chromosome numbers, biogeography, and divergence times. Journal of Systematics and Evolution, 54, 1–16.

46. Jarzembowski, E.A., Ross, A.J. 1996. Insect origination and extinction in the Phanerozoic. Geological Society London Special Publications, 102, 65–78.

47. Khramov, A.V. Bashkuev, A.S. and Lukashevich, E.D. 2020. The fossil record of long-proboscid nectarivorous insects. Entomological Review, 100, 881–968.

48. Khramov, A.V., Naugolnykh, S.V., Wegierek, P. 2022. Possible long-proboscid insect pollinators from the Early Permian of Russia. Current Biology, 32, 3815–3820.

49. Labandeira, C.C. 1998. Early history of arthropod and vascular plant associations. Annual Review of Earth and Planetary Sciences, 26, 329–377.

50. Labandeira, C.C. 2000. The paleobiology of pollination and its precursors, in Paleontological Society Papers, Vol. 6: Phanerozoic Terrestrial Ecosystems, Gastaldo, R.A. and DiMichele, W.A. (Eds.), 2000, p. 233.

51. Labandeira, C.C. 2014. Why did terrestrial insect diversity not increase during the Angiosperm Radiation? Mid-Mesozoic, plant-associated insect lineages harbor clues. In: Evolutionary Biology: Genome Evolution, Speciation, Coevolution and Origin of Life, Pierre Pontarotti (Ed.). Springer, pp. 261–299.

52. Labandeira, C.C., Eble, G.J. 2000. The fossil record of insect diversity and disparity. In: Anderson, J., Thackeray, F., Van Wyk, B. De Wit, M. (Eds.). Gondwana alive: biodiversity and the evolving biosphere. pp. 2–54. Witwatersrand Universit Press.

53. Labandeira CC, Sepkoski Jr JJ. 1993. Insect diversity in the fossil record. Science, 261, 310–315.

54. Labandeira, C.C., Kvaček, J., Mostovski, M.B. 2007. Pollination drops, pollen, and insect pollination of Mesozoic gymnosperms. Taxon, 56, 663–695.

55. Lavaut, E., Guillemim, M.-L., Colin, S., Faure, A., Coudret, J., Destombe, C., Valero, M. 2022. Pollinators of the sea: A discovery of animal-mediated fertilization in seaweed. Science, 377, 528–530.

56. Li, H.T., Yi, T.S., Gao, L.M., Ma, P., Zhang, T., Yang, J., Gitzendanner, M.A., Fritsch, P.W., Cai, J., Luo, Y., Wang, H., van der Bank, M., Zhang, S., Wang, Q., Wang, J., Zhang, Z., Fu, C., Yang, J., Hollingsworth, P.M., Chase, M.W., Soltis, D.E., Soltis, P., Li, D. 2019. Origin of angiosperms and the puzzle of the Jurassic gap. Nature Plants, 5, 461–470.

57. Lidgard, S., Crane, P.R. 1988. Quantitative analyses of the early angiosperm radiation. Nature, 331, 344–346.

58. Lloyd, G.T., Davis, K.E., Pisani, D., Tarver, J.E., Ruta, M., Sakamoto, M., Hone, D.W.E, Jennings, R., Benton, M.J. 2008. Dinosaurs and the Cretaceous Terrestrial Revolution. Proceedings of the Royal Society B: Biological Sciences, 275, 2483–2490.

59. Logomarsino, L.P., Condamine, F.L., Antonelli, A. Mulch, A., Davis, C.C. 2016. The abiotic and biotic drivers of rapid diversification in Andean bellflowers (Campanulaceae). New Phytologist, 210, 1430–1442.

60. Luo, C., Beutel, R.G., Engel, M. S., Liang, K., Li, L., Li, J., Xu, C., Vršanský, P., Jarzembowski, E.A., Wang, B. 2021. Life history and evolution of the enigmatic Cretaceous-Eocene Alienopteridae: A critical review. Earth-Science Reviews, 225, 103914.

61. Magallon S, Castillo A. 2009. Angiosperm diversification through time. American Journal of Botany 96, 349–65.

62. Magallón S, Hilu KW, Quandt D. 2013. Land plant evolutionary timeline: Gene effects are secondary to fossil constraints in relaxed clock estimation of age and substitution rates. American Journal of Botany, 100, 556–573.

63. Magallón, S., Gómez-Acevedo, S. Sánchez-Reyes, L.L., Hernández-Hernández, T. 2015. A metacalibrated time-tree documents the early rise of flowering plant phylogenetic diversity. New Phytologist, 207, 437–453.

64. Mayhew, P.J., Jenkins, G.B., Benton, T.G. 2008. A long-term association between global temperature and biodiversity, origination and extinction in the fossil record. Proceedings of the Royal Society B: Biological Sciences, 275, 47–53.

65. Mazet, N., Morlon, H., Fabre, P.-H., Condamine, F.L. 2022. Estimating clade-specific diversification rates and palaeodiversity dynamics from reconstructed phylogenies. bioRxiv preprint doi: https://doi.org/10.1101/2022.05.10.490920

66. McKenna, D.D. Sequeira, A.S. Marvaldi, A.E., Farrell B.D. 2009. Temporal lags and overlap in the diversification of weevils and flowering plants. Proceedings of the National Academy of Sciences of the United States of America, 106, 7083–7088.

67. Misof, B., Liu, S., Meusemann, K., Peters, R. S., Donath, A., Mayer, C., (et al.) & Zhou, X. (2014). Phylogenomics resolves the timing and pattern of insect evolution. Science, 346, 763–767.

68. Nel, A., Fleck, G., Garcia, G., Gomez, B., Ferchaud, P., Valentin, X. 2015. New dragonflies from the lower Cenomanian of France enlighten the timing of the odonatan turnover at the Early-Late Cretaceous boundary. Cretaceous Research, 52, 108–117.

69. Nepi, M., Little, S.A., Guarnieri, M., Nocentini, D., Prior, N.A., Gill, J., Tomlinson, P.B., Ickert-Bond, S.M., Pirone, C., Pacini, E. and von Aderkas, P. 2017. Phylogenetic and functional signals in gymnosperm ovular secretions. Annals of Botany, 120, 923–936.

70. Neubauer, T. A., Hauffe, T., Silvestro, D., Scotese, C. R., Stelbrink, B., Albrecht, C., Delicado, D., Harzhauser, M., Wilke, T. 2022. Drivers of diversification in freshwater gastropods vary over deep time. Proceedings of the Royal Society B: Biological Sciences, 289, 20212057.

71. Nicholson, D.B., Mayhew, P.J., Ross, A. J. 2015. Changes to the fossil record of insects through fifteen years of discovery. PLoS One, 10, e0128554.

72. Nie, Y., Foster, C.S.P., Zhu, T., Yao, R., Duchêne, D.A., Ho, S.Y.W., Zhong, B. 2020. Accounting for uncertainty in the evolutionary timescale of green plants through clock-partitioning and fossil calibration strategies. Systematic Biology, 69, 1–16.

73. Novikoff A., Barabasz-Krasny B. 2015. Modern plant systematics. General issues. Liga-Press, Lviv. – 686 p. (in Ukrainian).

74. Ollerton, J. 2017. Pollinator diversity: distribution, ecological function, and conservation. Annual Review of Ecology, Evolution, and Systematics, 48, 353–376.

75. Ollerton, J., Winfree, R., Tarrant, S. 2011. How many flowering plants are pollinated by animals? Oikos, 120, 321–326.

76. O’Meara, B.C., Smith, S.D., Armbruster, W.S., Harder, L.D., Hardy, C.R., Hileman, L.C., Hufford, L., Litt, A., Magallón, S., Smith, S.A., Stevens, P.F., Fenster, C.B. and Diggle, P.K. 2016. Non-equilibrium dynamics and floral trait interactions shape extant angiosperm diversity. Proceedings of the Royal Society B: Biological Sciences, 283, 20152304.

77. Onstein, R.E. 2020. Darwin’s second ‘abominable mystery’: trait flexibility as the innovation leading to angiosperm diversity. New Phytologist, 228, 1741–1747.

78. Peris, D., Davis, R.S., Engel, M.S., Delclòs, X. 2014. An evolutionary history embedded in amber: reflection of the Mesozoic shift in weevil-dominated (Coleoptera: Curculionoidea) faunas. Zoological Journal of the Linnean Society, 171, 534–553.

79. Peris, D., Pérez-de la Fuente, R., Peñalver, E., Delclòs, X., Barrón, E., Labandeira, C.C. 2017. False blister beetles and the expansion of gymnosperm-insect pollination modes before angiosperm dominance. Current Biology, 27, 897–904.

80. Peris, D., Labandeira, C.C., Barrón, E., Delclòs, X., Rust, J., Wang, B. 2020. Generalist pollen-feeding beetles during the mid-Cretaceous. iScience, 23, 100913.

81. Rainford, J.L., Hofreiter, M., Nicholson, D.B., Mayhew, P.J. 2014. Phylogenetic distribution of extant richness suggests metamorphosis is a key innovation driving diversification in insects. PLoS One, 9, e109085.

82. Ramírez-Barahona, S., Sauquet, H., Magallón, S. 2020. The delayed and geographically heterogeneous diversification of flowering plant families. Nature Ecology & Evolution, 4, 1232–1238.

83. Roddy, A.B., Théroux-Rancourt, G., Abbo, T., Benedetti, J.W., Brodersen, C.R., Castro, M., Castro, S., Gilbride, A.B., Jensen, B., Jiang, G.-F., Perkins, J.A., Perkins, S.D., Loureiro, J., Syed, Z., Thompson, R.A., Kuebbing, S.E., Simonin, K.A. 2020. The scaling of genome size and cell size limits maximum rates of photosynthesis with implications for ecological strategies. International Journal of Plant Sciences, 181, 75–87.

84. Rothwell, G.W., Crepet, W.L., Stockey, R.A. 2009. Is the anthophyte hypothesis alive and well? New evidence from the reproductive structures of Bennettitales. American Journal of Botany, 96, 296–322.

85. Sann, M., Niehuis, O., Peters, R., Mayer, C., Kozlov, A., Podsiadlowski, L., Bank, S., Meusemann, K., Misof, B., Bleidorn, C., Ohl, M. 2018. Phylogenomic analysis of Apoidea sheds new light on the sister group of bees. BMC Evolutionary Biology, 18, 1–15.

86. Sauquet, H., Von Balthazar, M., Magallón, S., Doyle, J.A., Endress, P.K., Bailes, E.J., et al., Schönenberger, J. 2017. The ancestral flower of angiosperms and its early diversification. Nature communications, 8, 16047.

87. Sauquet, H., Magallón, S. 2018. Key questions and challenges in angiosperm macroevolution. New Phytologist, 219, 1170–1187.

88. Sauquet, H., Ramírez-Barahona, S., Magallón, S. 2022. What is the age of flowering plants? Journal of Experimental Botany, erac130.

89. Schachat, S.R., Labandeira, C.L., Clapham, M.E., Payne, J.L. 2019. A Cretaceous peak in family-level insect diversity estimated with mark–recapture methodology. Proceedings of the Royal Society B: Biological Sciences, 286, 20192054.

90. Silvestro, D., Bacon, C.D., Ding, W., Zhang, Q., Donoghue, P.C.J., Antonelli, A., Xing, Y. 2021. Fossil data support a pre-Cretaceous origin of flowering plants. Nature Ecology and Evolution, 5, 449–457.

91. Silvestro, D., Cascales-Miñana, B., Bacon, C.D., Antonelli, A. 2015. Revisiting the origin and diversification of vascular plants through a comprehensive Bayesian analysis of the fossil record. New Phytologist, 207, 425–436.

92. Silvestro, D., Salamin, N., Antonelli, A., Meyer, X. 2019. Improved estimation of macroevolutionary rates from fossil data using a Bayesian framework. Paleobiology, 45, 546–570.

93. Simonin, K.A., Roddy, A.B. 2018. Genome downsizing, physiological novelty, and the global dominance of flowering plants. PLoS Biology, 16, e2003706.

94. Simpson, B.B., Neff, J.L. 1981. Floral rewards: Alternatives to pollen and nectar. Annals of the Missouri Botanical Garden, 68, 301–322.

95. Sokoloff, D.D., Remizowa, M.V., El, E.S., Rudall, P.J., Bateman, R.M. 2020. Supposed Jurassic angiosperms lack pentamery, an important angiosperm-specific feature. New Phytologist, 228, 420–426.

96. Soltis, P.S., Folk, R.A., Soltis, D.E. 2019. Darwin review: angiosperms phylogeny and evolutionary radiations. Proceedings of the Royal Society B: Biological Sciences, 286, 20190099.

97. Song, H., Béthoux, O., Shin, S., Donath, A., Letsch, H., Liu, S., McKenna, D.D., Mang, G., Misof, B., Podsiadlowski, L., Zhou, X., Wipfler, B., Simon, S. 2020. Phylogenomic analysis sheds light on the evolutionary pathways towards acoustic communication in Orthoptera. Nature Communications, 11, 4939.

98. Suchan, T., Alvarez, N. 2015. Fifty years after Ehrlich and Raven, is there support for plant–insect coevolution as a major driver of species diversification? Society Entomologia Experimentalis et Applicata, 157, 98–112.

99. Toon, A., Terry, L.I., Tang, W., Walter, G.H., Cook, L.G. 2020. Insect pollination of cycads. Austral Ecology, 43, 1033–1058.

100. Vamosi, J.C., Magallón, S., Mayrose, I., Otto, S.P., Sauquet, H. 2018. Macroevolutionary patterns of flowering plant speciation and extinction. Annual Review of Plant Biology, 69, 1–22.

101. Vamosi, J.C., Vamosi, S.M. 2011. Factors influencing diversification in angiosperms: at the crossroads of intrinsic and extrinsic traits. American Journal of Botany, 98, 460–471.

102. Van del Kooi, C., Ollerton, J. 2020. The origins of flowering plants and pollinators. Science, 368, 1306–1308.

103. Van der Niet, T., Peakall, R., Johnson, S.D. 2014. Pollinator-driven ecological speciation in plants: new evidence and future perspectives. Annals of Botany, 113, 199–211.

104. Veizer, J., Prokoph, A. 2015. Temperatures and oxygen isotopic composition of Phanerozoic oceans. Earth-Science Reviews, 146, 92–104.

105. Wardhaugh, C.W. 2015. How many species of arthropods visit flowers? Arthropod-Plant Interactions, 9, 547–565.

106. Waser, N.M. Chittka, L., Price, M.V., Williams, N.M., Ollerton, J. 1996. Generalization in pollination systems, and why it matters. Ecology, 77, 1043–1060.

107. Westerhold, T., Marwan, N., Drury, A. J., Liebrand, D., Agnini, C., Anagnostou, E., Barnet, J.K., Bohaty, S.M., de Vleeschouwer, D., Florindo, F., Frederichs, T., Hodell, D.A., Holbourn, A.E., Kroon, D., Lauretano, V., Litter, K., Lourens, L.J., Lyle, M., Pälike, H., Röhl, U., Tian, J., Wilkens, R., Wilson, P.A., Zachos, J.C. (2020). An astronomically dated record of Earth’s climate and its predictability over the last 66 million years. Science, 369, 1383–1387.

108. Wiegmann, B.M., Trautwein, M.D., Winkler, I.S., Barr, N.B., Kim, J.-W., Lambkin, C., Bertone, M.A., Cassel, B.K., Bayless, K.M., Heimberg, A.M., Wheeler, B.M., Peterson, K.J., Pape, T., Sinclair, B.J., Skevington, J.H., Blagoderov, V., Caravas, J., Kutty, S.N., Schmidt-Ott, U., Kampmeier, G.E., Thompson, F.C., Grimaldi, D.A., Beckenbach, A.T., Courtney, G.W., Friedrich, M., Meier, R., Yeates, D.K. 2011. Episodic radiations in the fly tree of life. Proceedings of the National Academy of Sciences of the United States of America, 108, 5690–5695.

